# Expression diversity of cichlid MHC alleles

**DOI:** 10.1101/2024.01.23.576710

**Authors:** Seraina E. Bracamonte, Pascal I. Hablützel, Carlos Lozano-Martín, Marta Barluenga

**Affiliations:** Museo Nacional de Ciencias Naturales - CSIC, Madrid, Spain; Flanders Marine Institute, Oostende, Belgium; Vrije Universiteit Brussel, Brussels, Belgium

**Keywords:** Major histocompatibility complex, gene expression, cichlids, host-parasite interactions

## Abstract

Diversification by natural selection is a major source of biodiversity. The selective pressures imposed by parasites on their hosts have the potential to increase diversity at immunologically relevant genes and maintain high polymorphism within and among populations. The immune genes of the major histocompatibility complex (MHC) are a hallmark of parasite-mediated selection. These genes frequently show local adaptation to the prevailing parasites. Although associations between parasites and alleles have been identified, they are often imperfect, indicating that parasite-mediated selection goes beyond sequence identity. Here, we explored allele-specific expression of MHC class IIB in cichlid fish and its relationship with sequence diversity. This revealed a few highly abundant alleles with consistently low expression, and many relatively rare alleles with high expression levels that were variable among individuals. Alleles that formed functional supertypes also had similar expression. This suggests that rare alleles may be those responding to parasites and that similar functionality may be provided by different alleles in different individuals. Our results further suggest that there could be an optimal number of MHC alleles per individual and that deviations from this optimum are compensated by adjusting expression levels. The common and lowly expressed alleles may have attained different functions, a patterns that appears to be consistent in Neotropical and African cichlids. Our study shows that MHC expression is highly variable among alleles potentially interacting with parasites, providing an additional substrate for selection. Furthermore, our study exemplifies how combining genetic diversity with detailed expression information can help identifying functionally relevant diversity.

## Introduction

Host-parasite interactions are a major force driving evolution at different levels of biological organisation and in particular can lead to an excess of genetic diversity in some genomic regions (Betts et al., 2018; Ebert & Fields, 2020; Schmid-Hempel, 2013; Summers et al., 2003). Infection by parasites triggers the evolution of defence responses in the host that try to mitigate the costs of infection, which in turn induces parasites to adapt to the new conditions, culminating in an evolutionary arms race (Haldane, 1949; Woolhouse et al., 2002). These highly dynamic interactions generate and maintain genetic diversity among and within populations. A few genomic regions coding for genes of the host’s immune system show evidence of parasite-mediated balancing selection and consequently maintain remarkably high levels of polymorphism (Dexter et al., 2023; Eizaguirre & Lenz, 2010). This is particularly true for the genes of the major histocompatibility complex (MHC). The MHC is at the core of the immune defence of vertebrates against parasites. MHC molecules present foreign antigens to T cells which initiates an adaptive immune response that is specific to the infecting agent (Murphy & Weaver, 2017; Penn & Ilmonen, 2005). There are two classes of MHC molecules that are encoded by highly polymorphic gene families. MHC class I is expressed on most body cells and binds peptides from intracellular parasites and pathogens (e.g. viruses). MHC class II is only expressed on specialized antigen-presenting cells and binds peptides from extracellular parasites (e.g. metazoans; Murphy & Weaver, 2017; Penn & Ilmonen, 2005).

MHC class I and class II are highly dynamic multigene families, whose evolution are characterized by birth-and-death processes, where new genes are created by duplications, some duplicated genes are retained in the genome, and pseudogenization and neo-functionalization are common features (Klein et al., 1998; Nei & Rooney, 2005). Positive selection on the sites interacting directly with the parasitic antigens facilitates the accumulation of mutations in the different paralogous gene copies. This process leads to high polymorphism and copy number variation among and within species and populations. As a result, each individual, population, and species, displays a unique set of multiple variants of MHC class I and class II molecules, which allows responding to a diverse array of parasites. The extreme levels of polymorphism are then maintained by parasite-mediated balancing selection, with MHC allele pools adapted to the local parasite communities (Bernatchez & Landry, 2003; Eizaguirre & Lenz, 2010; Spurgin & Richardson, 2010). Locally distinct allele pools were inferred for a wide range of vertebrates (Cammen et al., 2011; Kyle et al., 2014; Larson et al., 2014; Miller et al., 2010; Minias et al., 2023; Talarico et al., 2019) which tracks divergence of infecting parasite communities (Eizaguirre et al., 2012b; Pavey et al., 2013; Peng et al., 2021; Stutz & Bolnick, 2017). Additionally, specific alleles are associated with resistance or susceptibility to specific parasites (Bonneaud et al., 2006; Eizaguirre et al., 2012a; Peng et al., 2021). Associations with parasite infections were also described for groups of functionally similar alleles, i.e. supertypes, rather than single alleles (Schmid et al., 2023; Sepil et al., 2013). These associations are, however, often imperfect and for the vast majority of alleles no associations with parasite infections could be identified, suggesting that parasite-mediated selection may not act solely at the MHC sequence level.

The immune response to parasites may be modulated by the regulation of expression levels of particular MHC alleles (Handunnetthi et al., 2010). Different studies have suggested that not all MHC alleles are expressed (Bracamonte et al., 2015; Gaczorek et al., 2023; Hofmann et al., 2017; Sin et al., 2012), and that expression is tissue-specific (Gerdol et al., 2019; Harstad et al., 2008). However, less focus has been placed on the importance of MHC regulation for the response to parasites. Studies that focussed on the expression of specific MHC class IIB alleles or on overall expression found that alleles and genotypes differ in expression (Harstad et al., 2008; Schwensow et al., 2019), expression of alleles or genotypes increases in the presence of specific parasites (Axtner & Sommer, 2012), and increased expression may compensate for suboptimal individual MHC diversity (Wegner et al., 2006). Despite that, it remains unresolved whether MHC expression levels correlate with parasite infections. Identifying such associations requires higher, allele-specific resolution of MHC expression. Studies that thoroughly assess allele-specific expression are scarce, particularly for non-model organisms, and focusses predominantly on MHC class I. In birds, large variation in expression among MHC class I alleles were identified (Drews et al., 2017; Drews & Westerdahl, 2019). In these studies, a set of alleles showed consistently low expression, and they were attributed to the non-classical MHC. The non-classical MHC is not involved in antigenic peptide-binding, but serves different functions in the immune response, from assisting with peptide-binding to binding parasitic lipid molecules (Mak et al., 2014; Rodgers & Cook, 2005).

It is characterized by low levels of polymorphism and expression. Classical MHC class I alleles, which are highly polymorphic, have been shown to have pronounced variation in expression (Chappell et al., 2015; Drews et al., 2017; Kaufman, 2018). For MHC class II, allele-specific high and low expression was identified in humans and mice with consequences for disease risk, although environmental factors may have a considerable effect on expression levels (Handunnetthi et al., 2010). Similarly, a study on newts inferring allele expression from whole transcriptome sequencing identified several highly expressed alleles and a strongly supported cluster of alleles with practically no expression (Gaczorek et al., 2023), suggesting that expression of MHC class II may follow a similar pattern as expression of MHC class I. In this study, we used both allele-specific sequencing of the MHC IIB and transcriptome-wide expression data for a comparative exploration of expression variation in this gene family across cichlid fish.

Cichlid fish are one of the most species-rich vertebrate family (Salzburger, 2018; Seehausen & Wagner, 2014). They inhabit tropical and subtropical freshwater systems where they have rapidly diversified as a result of natural and sexual selection (Barluenga & Meyer, 2010; Kautt et al., 2020; Matschiner et al., 2020; Ronco et al., 2021; Salzburger, 2009). Exposure to divergent parasite communities could be a mechanism contributing to cichlid diversification (Blais et al., 2007; Hablützel et al., 2017; Karvonen et al., 2018; Maan et al., 2008). Cichlid fish have one of the largest MHC class II repertoires described so far (Hofmann et al., 2017; Málaga-Trillo et al., 1998), which could be driven by parasite mediated selection. MHC allele pools exhibit considerable differentiation between populations and closely related species (Blais et al., 2007; Bracamonte et al., 2022; Hablützel et al., 2016), and at the same time, allelic lineages are shared among independent radiations even across continents (Hablützel et al., 2013; Lozano-Martín et al., 2023). Similar to other systems, MHC allele pool divergence in cichlid fish did not entirely correspond to divergence in parasite communities (Hablützel et al., 2016; Meyer et al., 2019), indicating that selection may act at different levels on the MHC, including expression.

In this study, we aimed to understand the relationship between MHC IIB allele repertoires and expression patterns in the Neotropical Midas cichlid fish *Amphilophus* spp. We investigated whether frequent and ubiquitous alleles are the key to resistance and survival with higher levels of expression, or, on the contrary, whether rare alleles may be those responding to parasite infection pressures and showing higher levels of expression. To test for this, we identified MHC class IIB alleles in populations along the species distribution, determined frequencies and mean expression, and classified them into *highly expressed* and *lowly expressed* alleles. We grouped functionally similar alleles into supertypes, and measured their frequency and expression. We evaluated whether expression patterns are homogeneous within supertypes or not. To determine whether expression variation is a consistent pattern of cichlid MHC class II, we further identified expression levels of putative MHC class II loci in several African cichlid species. Lastly, we evaluated if expression and number of alleles per individual are correlated.

## Material and methods

### Sampling

We collected samples of Midas cichlid fish in Nicaragua in December in 2017 and 2018 from the two Great Nicaraguan lakes and five crater lakes (Fig. 1). We caught fish with gill nets, and euthanized them with an overdose of Tricaine mesylate (MS-222) on cold water, following the procedures approved by the American Veterinary Medical Association Guidelines for Euthanasia of Animals: 2020 edition (available at https://www.avma.org/sites/default/files/2020-02/Guidelines-on-Euthanasia-2020.pdf). We stored muscle tissue in absolute ethanol at −20 °C and we collected the spleen in RNA*later* Stabilization Solution (Ambion, USA) and stored it at −80 °C thereafter.

**Figure 1.**
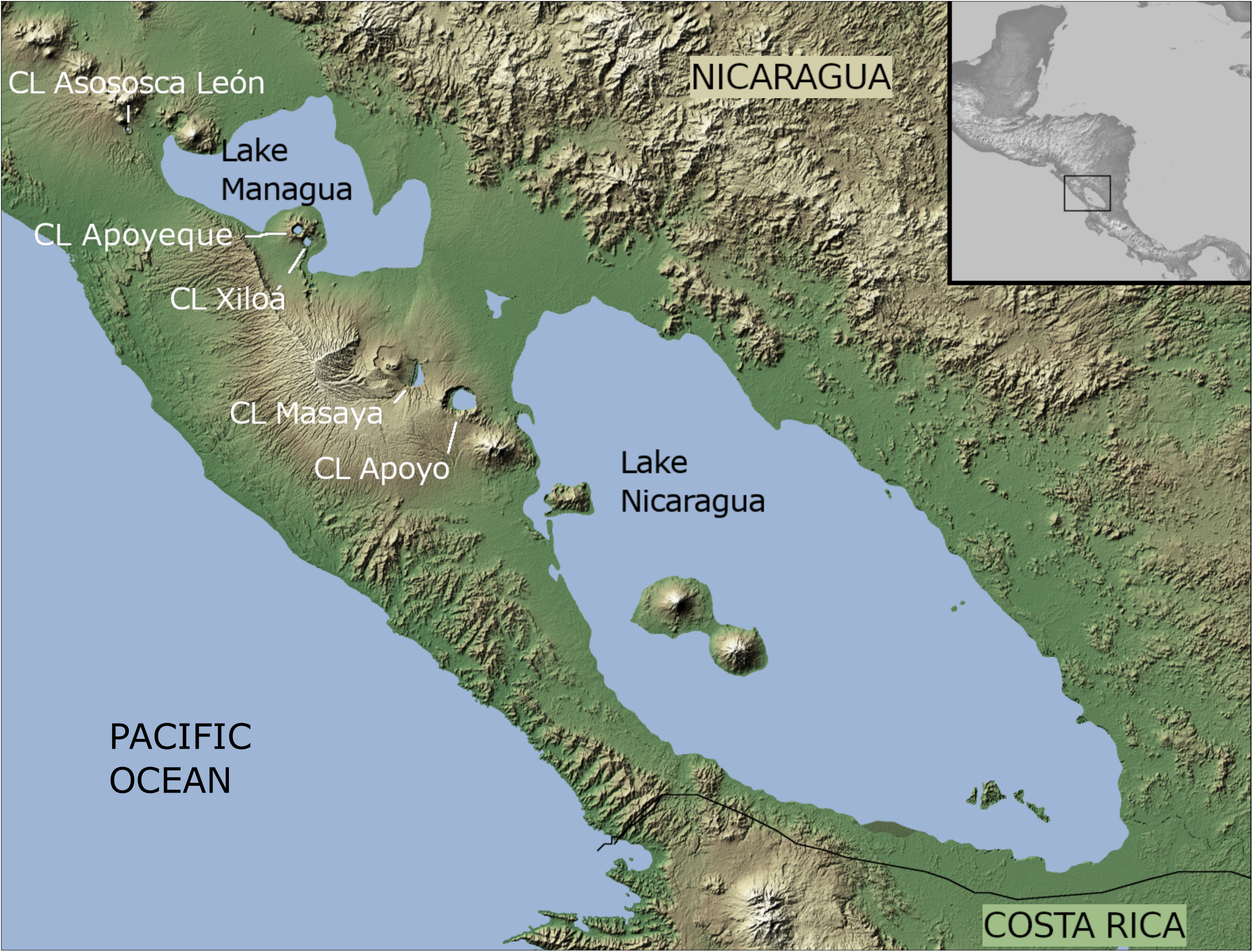
Map of the great lakes region in Nicaragua (modified from NASA Photojournal PIA03364). Sampled lakes are indicated. CL = Crater Lake.

### DNA and RNA extraction

We extracted DNA from fin, muscle, or spleen with the DNeasy Blood and Tissue kit (Qiagen, Germany) following the manufacturer’s recommendations. We incubated a piece of tissue with 20 µl of Proteinase K in 180 µl of ATL buffer at 56 °C for approximately 3 hours. We mixed the lysate with 200 µl of AL buffer and 200 µl of ethanol and transferred it to the spin column. We cleaned the DNA first with 500 µl of AW1 buffer and then with 500 µl of AW2 buffer and eluted it in 200 µl of DNase/RNAse-free water. We removed remnant RNA by incubating the DNA with 14 U of RNase A (Qiagen, Germany) at room temperature for 4 min and inactivated the RNase by freezing at −20 °C. DNA was quantified with Qubit (Invitrogen, USA).

We extracted RNA from a fragment of 5 x 5 mm of spleen, which we homogenized with ceramic beads in 250 μl of TRI Reagent at 6800 rpm for 3 x 20 s in a Precellys Evolution homogenizer (Bertin Instruments, France). We incubated the homogenate for 5 min at room temperature, then added 50 μl of chloroform. We incubated the mixture for 2 min at room temperature and centrifuged it for 15 min at 12500 rpm at 4 °C. We transferred the supernatant to a new tube, added 125 μl of isopropanol, and incubated it overnight at −80 °C. We transferred the liquid to a Direct-zol RNA Miniprep (Zymogen, USA) spin column and cleaned the RNA following the manufacturer’s instructions. In short, we washed the spin column twice with 400 μl of RNA PreWash and once with 700 μl of RNA Wash Buffer. We eluted the RNA in 40 μl of DNase/RNase-free water and treated it with DNase I (Thermo Scientific, USA) following the manufacturer’s recommendations. RNA quantity and quality were measured with a NanoDrop 1000 (Thermo Scientific, USA).

#### cDNA synthesis and qPCR testing

We synthesized cDNA with the RevertAid First Strand cDNA Synthesis Kit (Thermo Scientific) following the manufacturer’s instructions for gene-specific primers. Gene-specific reverse transcription primers targeted the MHC IIB (5’-AACATGGCAGGATG-3’) and two housekeeping genes (ACTB 5’-TCCCTGTTGGCTTT-3’, EF1A 5’-GTCGTTCTTGCTGT-3’). We mixed approximately 450 ng of DNase-treated RNA with 20 pmol of each reverse transcription primer, 4 μl of 5x reaction buffer, 1 μl of RiboLock RNase Inhibitor, 2 μl of dNTPs, and 1 μl of RevertAid M-MuLV reverse transcriptase in a final volume of 20 μl and incubated it for 60 min at 42 °C, then for 5 min at 70 °C, and stored it at −80 °C.

We diluted the cDNA 1:20 and amplified MHC IIB, ACTB, and EF1A in a single PCR reaction. We reduced the amplification of the housekeeping genes with blocking reverse primers with a 3’ amino linker. We determined the optimal blocking primer concentrations with qPCRs, gradually increasing the concentration of blocking reverse primers and decreasing the concentration of normal reverse primers, keeping the total concentration of reverse primers constant at 0.3 μM. We approximated the amount of PCR product with the height of peaks of the melting curves corresponding to each gene. We deemed the concentration of blocking primers that rendered peaks of similar heights as optimal. We prepared qPCR reactions as described below for regular PCRs but using the FastStart Universal SYBR Green Master (ROX). Cycling conditions were as follows: initial denaturation of 10 min at 95 °C and 40 cycle of 95 °C for 15 s, 50 °C for 30 s, and 72 °C for 1 min. This was followed by the melting analysis. We determined the optimal number of thermal cycles for regular PCRs from the amplification curves in order to avoid reaching the plateau phase.

#### MHC amplification and sequencing

MHC IIB was amplified from DNA with one forward and two reverse primers (Bracamonte et al., 2022) and sequenced on a Illumina MiSeq by LGC Genomics (Germany). Twenty-one samples were amplified and sequenced in duplicates. Briefly, PCRs were done in 20 µl containing 15 pmol of each primer and 1.5 U of MyTaq polymerase. Cycling conditions were as follows: 2 min denaturation at 96 °C followed by 20 cycles of 96 °C for 15 s, 50 °C for 30 s, 70 °C for 1 min. Following primer removal, libraries were prepared using the MiSeq v3 chemistry. Paired-end sequencing resulted in a mean of 15700 2x300 bp reads per sample.

We amplified MHC IIB and housekeeping genes from cDNA in a single reaction for each sample. We did PCRs in a reaction volume of 25 μl containing 12.5 μl 2x FastStart PCR master mix (Roche, Germany), one forward and two reverse MHC IIB primers (Bracamonte et al., 2022), one forward, one reverse, and one blocking reverse primer for each housekeeping gene, and 6.25 μl of diluted cDNA. Primer sequences and final concentrations are given in Table 1. Cycling conditions were the same as for qPCRs with only 23 cycles followed by a final elongation step of 7 min at 72 °C. We checked amplification success on an agarose gel. Adapter ligation and paired-end sequencing on an Illumina NextSeq 500/550 v2 were done at LGC Genomics (Germany) using 20 μl PCR product as starting material. This resulted in a mean of 636677 2x150 bp reads per sample.

**Table 1.**
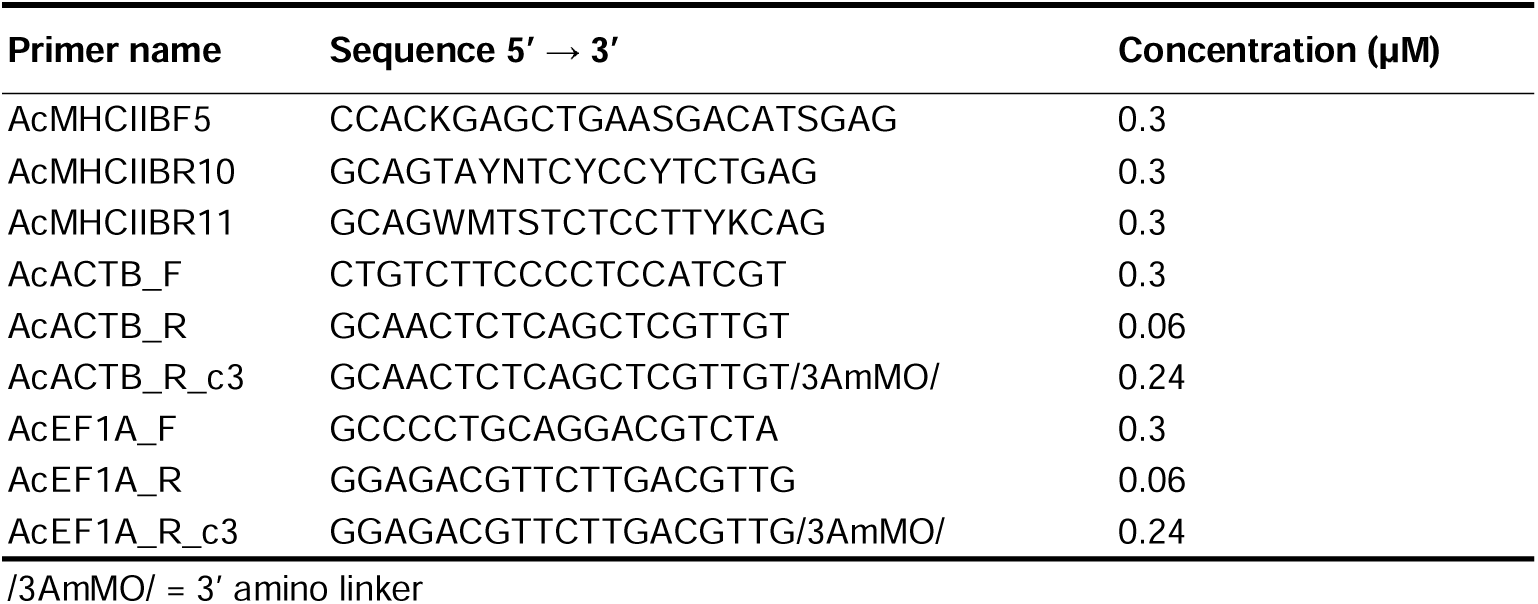
Primer sequences and concentrations used in PCR reactions. Each PCR contained all primers at indicated concentrations.

### Read processing

We demultiplexed reads of both DNA and cDNA sequencing and trimmed adapters with bcl2fastq v2.20 (Illumina, 2017) allowing for up to two mismatches in the barcode and discarding trimmed reads shorter than 20 base pairs (bp). We assembled DNA reads into alleles using the AmpliSAT pipeline (Sebastian et al., 2016). We merged reads with AmpliMERGE and cleaned them with AmpliCLEAN, discarding reads with mean Phred scores < 30 and missing forward or reverse primers. We trimmed primers with a custom python script before inferring allele profiles for each sample with AmpliSAS. To determine the best per amplicon frequency (PAF) filter, we ran AmpliSAS with values from 2 to 3.5 with incremental steps of 0.1. We compared the number of inferred alleles per sample between consecutive PAF values. The PAF value for which the number of alleles per sample differed least from the less stringent value and from the more stringent value was considered optimal. Additionally, we compared the agreement of samples that were PCR-amplified and sequenced in duplicates for each PAF. Both methods identified 2.7 as the best PAF for our data and all further analyses used the alleles called with this filter. Additional AmpliSAS parameters were as described in Bracamonte et al. (2022). Alleles could not be called for three samples due to the low number of reads that passed quality filtering. We renamed newly identified alleles in accordance with the naming system that we introduced in this previous study (“acXXX” where XXX follows consecutive numbering). We removed an allele with a 1-bp deletion leading to a premature stop codon from further analyses.

We trimmed primers from cDNA reads with a custom python script and merged them with BBMerge v38.90 (Bushnell et al., 2017) using the maxstrict setting and requiring a match between ratio mode and flat mode. We discarded reads shorter than 132 bp (expected length MHC -10 bp) and longer than 178 bp (expected length EF1A +10 bp) using the script reformat.sh from BBMap v38.90. We mapped the surviving reads of each sample to the corresponding MHC profile, which included the sequences of the housekeeping genes ACTB and EF1A, with Bowtie2 v2.4.4 in end-to-end alignment mode (Langmead & Salzberg, 2012). Seed length for multiseed alignment was set to 32 bases with a seed interval of 1 base. Re-seeding was set to 3 allowing for up to 20 seed extension attempts. Maximum and minimum mismatch penalties were 12 and 6, respectively, and read gap opening and extension penalties were 8 and 5. We sorted SAM files by coordinate and converted them to BAM files with samtools v1.9 (Li et al., 2009). We removed reads with mapping quality < 8. We quantified reads per MHC allele and housekeeping gene for each sample separately with featureCounts (Liao et al., 2014) from the R package Rsubread v2.4.2 (Liao et al., 2019) using the MHC allele profile of the respective sample as reference. We could not map and quantify reads for one sample, because its MHC profile could not be inferred from DNA, and we additionally removed 18 samples due to low read numbers from RNA (fewer than 10% of the mean number of reads).

#### Quantification of African cichlid MHC expression

We used published data sets of transcriptome sequencing of the gills of adult individuals of several cichlid species from Lake Tanganyika (El Taher et al., 2020; BioProject PRJNA552202; species, sex, and accession numbers are given in table S1). We mapped the reads separately to the *Oreochromis niloticus* reference genome and to a curated list of representative MHC alleles of each locus defined in Hablützel et al (2013) with HISAT2 v2.2.1 (Kim et al., 2019) and quantified the reads with HTSeq (Putri et al., 2022). We combined these two sets of read counts for further analyses.

### Data analysis

We constructed a maximum likelihood phylogenetic tree of MHC IIB with the *Amphilophus* and the African alleles with IQ-Tree v2.0.6 (Minh et al., 2020). We determined the best-fit nucleotide substitution model with ModelFinder (Kalyaanamoorthy et al., 2017) implemented in IQ-Tree using separate partitions for each codon position. This identified model TVM+F+G4 for positions 1 and 2 and model K3P+G4 for position 3, and we inferred the phylogeny using these two partitions (Chernomor et al., 2016). We obtained node support with the ultrafast bootstrap method (Hoang et al., 2018) with 10,000 replications.

### Midas cichlid MHC

To group alleles into supertypes, we identified positively selected sites for all known *Amphilophus* MHC class IIB alleles with four different methods: FEL, FUBAR (all available on the datamonkey server; Pond et al., 2020; Weaver et al., 2018), and the two model comparisons implemented in CodeML (models M2a *vs*. M1a and M8 *vs*. M7; Yang, 2007). We considered a site as positively selected if it was identified by at least two methods. We inferred the supertype of each allele as described in Bracamonte et al (2022). We calculated nucleotide diversity (π), the ratio of non-synonymous substitutions per non-synonymous site (dN) over synonymous substitutions per synonymous site (dS), and gene-wide positive selection for each supertype with MEGA11 (Tamura et al., 2021).

For all further analyses, we only included individuals for which we had both the MHC profile and expression estimates. We did all data analyses in R v4.2.1 (R Core Team, 2022), unless otherwise indicated. We normalized expression of each MHC allele per sample relative to the housekeeping genes. For this, we divided the number of reads of each MHC allele by the number of reads of each housekeeping gene and considered the mean of these normalizations as relative expression. We calculated frequency of occurrence and mean expression for each allele and supertype and determined the distribution of allele expression among supertypes. We classified alleles into *highly expressed* alleles and *lowly expressed* alleles. For this, we sorted the alleles by mean expression and calculated the change of expression from one allele to the next lower expressed allele. We set the threshold between *highly expressed* and *lowly expressed* alleles where the change of expression exceeded half the expression of the higher expressed allele for the first time. We considered alleles with higher expression as *highly expressed* and alleles with lower expression as *lowly expressed*. The latter category also included the non-expressed alleles. We mapped mean expression and supertype identity to the allele phylogeny with ggtree (Yu et al., 2017). For supertypes, we determined the relationship between π and mean relative expression with a Spearman correlation.

For each individual, we determined the number of alleles, the number of expressed alleles, and mean relative expression. We also determined these parameters for *highly expressed* and *lowly expressed* alleles separately. We examined the relationship among these variables with Pearson correlations. To test whether expression may compensate for suboptimal numbers of alleles, we correlated mean expression with the absolute difference of the number of alleles from the median number of alleles. Furthermore, we identified the distribution of supertypes among individuals.

### African cichlid MHC

We standardized read counts by library size, estimated dispersion, and normalized counts with DESeq2 v1.36.0 (Love et al., 2014). We extracted normalized read counts of MHC IIB and MHC IIA alleles and used them as proxy for expression. We calculated median expression of each allele and tested for differential expression between them using Kruskal-Wallis tests and pairwise Mann-Whitney U tests with false discovery rate correction of p-values.

## Results

We found a total of 154 alleles in 309 Midas cichlid samples. One allele (Amci-DXB*040101) was present in all individuals, and another (ac001) was present in all but three individuals, while 52 were singletons, *i.e*. they occurred in a single individual (Table S2). In total, 143 alleles were expressed in at least one individual, while 11 were not found expressed in any individual (Table S2). For the non-expressed alleles, seven were singletons, three were rare (present in only a few individuals), while one was relatively abundant (present in 16.5% of all individuals; Fig. 2a, b). For the expressed alleles, 18 were only expressed in some of the individuals in which they occurred covering the entire range of expression, while the remaining 125 expressed alleles were always expressed (Table S2). Twenty-three of these alleles were classified as *lowly expressed* according to our criteria. This included the 11 non-expressed alleles and eight alleles that were expressed only in some of their carriers. The ubiquitous allele Amci-DXB*040101 was also *lowly expressed*. Mean relative expression of the expressed alleles ranged from 4*10^-5^ to 24.2 (*lowly expressed*: 0-0.07, *highly expressed*: 0.16-24.2; Table S2).

**Figure 2.**
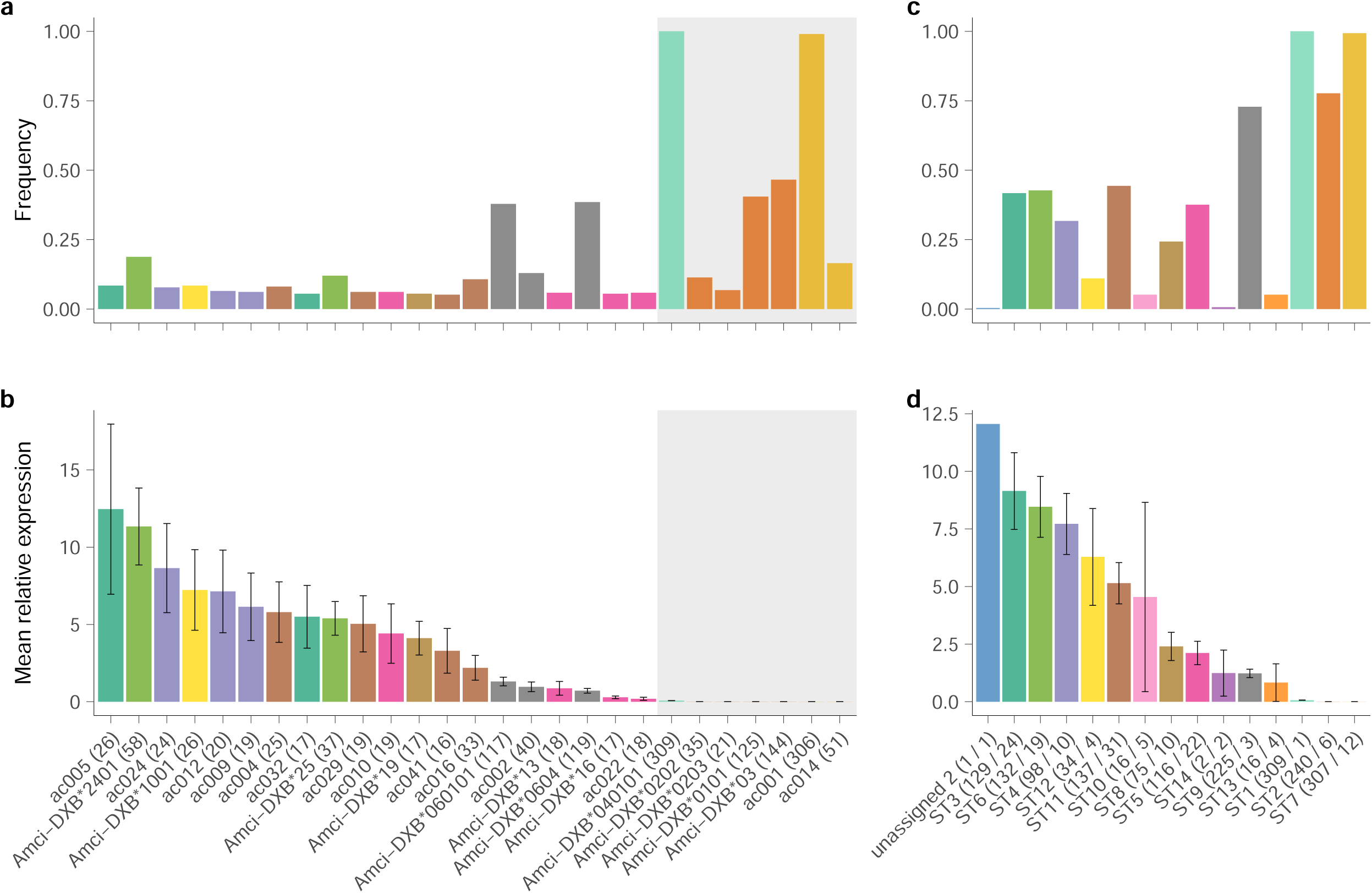
Frequency (a) and mean relative expression (b) of the 27 most frequent MHC IIB alleles (i.e. present in > 5% of all individuals), and frequency (c) and mean relative expression (d) of the 13 supertypes. Alleles are coloured by supertype. *Lowly expressed* alleles are shaded in grey.

Midas cichlid MHC alleles clustered into 14 supertypes (ST), with four to 38 alleles per supertype, and two unassigned alleles that did not cluster in any supertype (Fig. 3, Table S3). Sequence diversity (π) ranged from 0.019 for ST9 to 0.084 for ST10. Mean relative expression of supertypes ranged from 0.0002 for ST7 to 9.1 for ST3 (Table S3). Supertype size was not significantly correlated with π (r = 0.43, p = 0.12), this was particularly evident when considering only similarly sized supertypes. Mean relative expression was also not correlated with supertype size (r = 0.25, p = 0.36).

**Figure 3.**
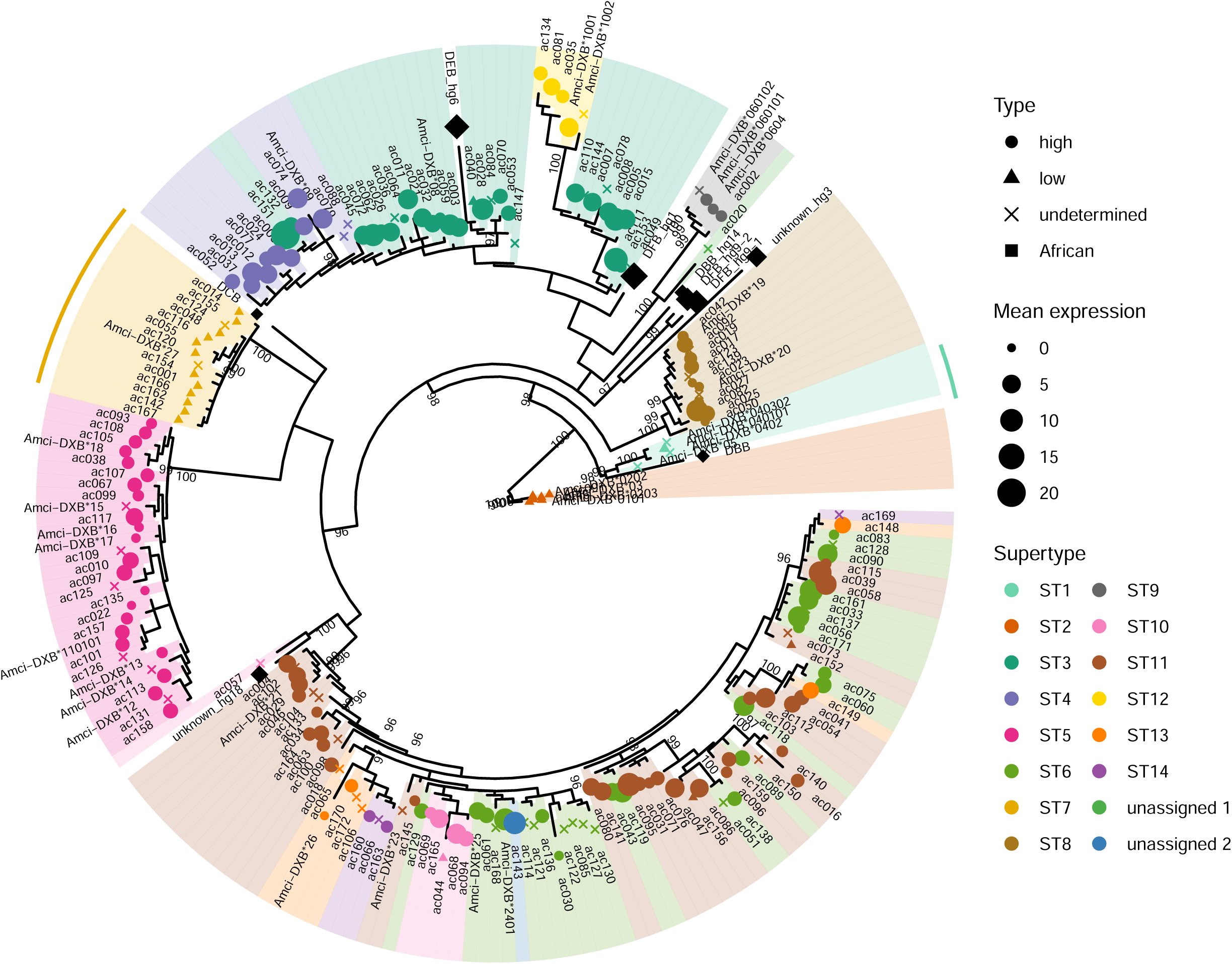
Phylogeny of African and Neotropical MHC IIB alleles. Symbols (type) indicate if alleles are highly expressed, lowly expressed, undetermined (alleles without expression information), or of African origin. The size of the symbol shows mean expression. Colours refer to supertype for Midas cichlid alleles, African loci are shown in black. The two African alleles with lowest expression cluster with the lowly expressed Midas supertypes ST1 and ST7, indicated by bars. Since expression for Midas and African alleles was measured at different scales, read counts of African alleles were divided by 100 to match the scale of Midas allele expression.

Supertypes ST1, ST2, and ST7, which only contained *lowly expressed* alleles, formed three well-defined clusters on the MHC phylogeny (Fig. 3) with relatively low sequence diversity (π_ST1_ = 0.032, π_ST2_ = 0.026, π_ST7_ = 0.022, mean π_remaining_ = 0.056; Fig. 4, Table S3). Despite that, ST7 had one of the highest dN/dS ratios and was under gene-wide positive selection. Alleles of ST5 and ST8 also formed two well-defined clusters. The alleles of these supertypes all belonged to the highly expressed group and showed moderate expression (mean_ST5_ = 2.1, mean_ST8_ = 2.4) and moderate sequence diversity (π_ST5_ = 0.048, π_ST8_ = 0.033; Fig. 3, Table S3). Mean relative expression of supertypes was significantly positively correlated with π (r = 0.6, p = 0.02; Fig. 4).

**Figure 4.**
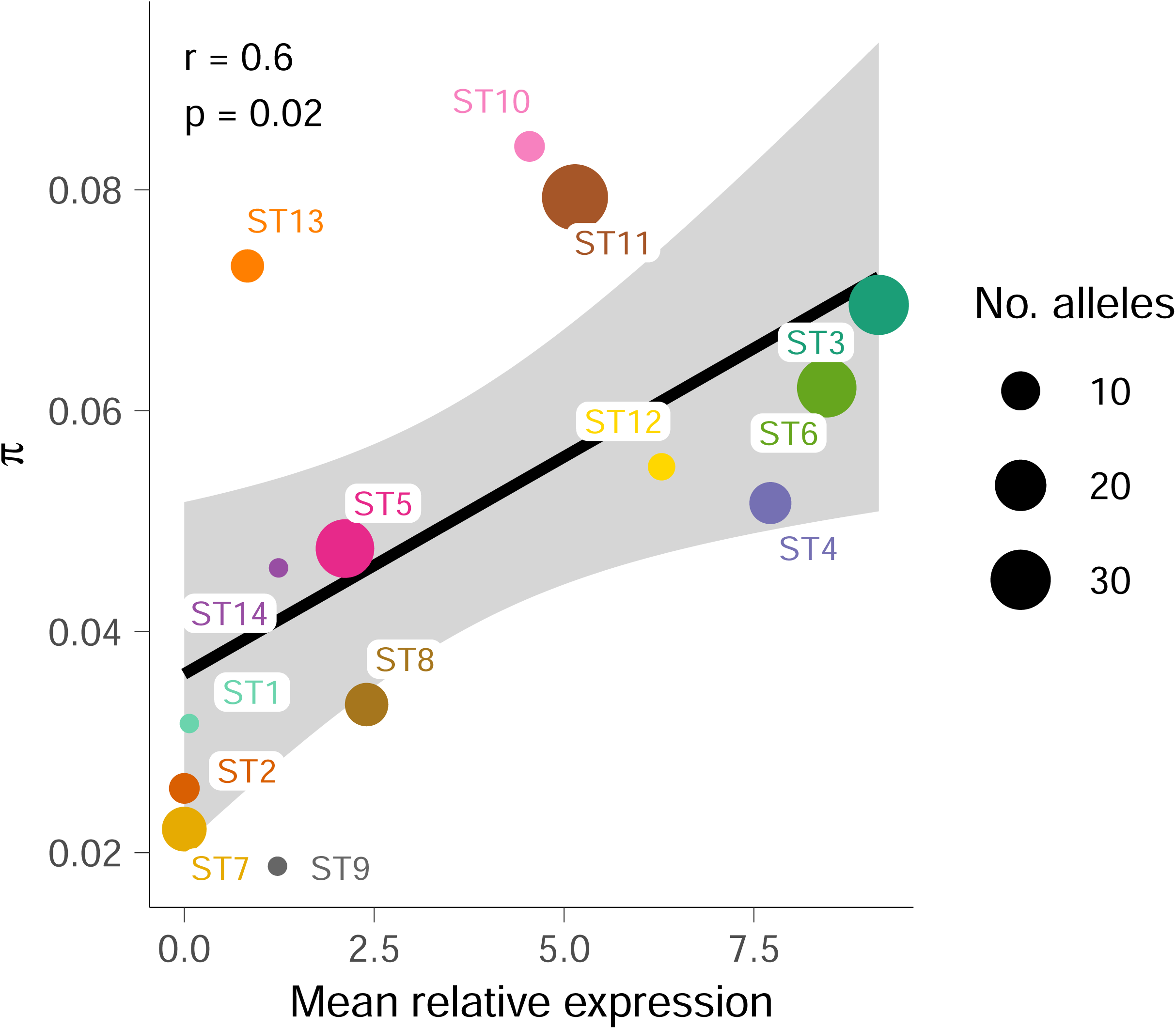
Correlation between supertype mean expression and sequence diversity (π). The 95% confidence interval is shaded in grey. The size of the circles corresponds to the number of alleles that of each supertype (range: 4-38).

Twenty-seven alleles occurred in more than 5% of all individuals, and we considered these common alleles (see Bracamonte et al., 2022). Of these common alleles, seven were lowly expressed. Despite their low expression, they were among the most frequent alleles with two occurring in > 99% of all individuals (Fig. 2a, b). In fact, the four most frequent alleles belonged to the *lowly expressed* group. Among the *highly expressed* alleles, the two most frequent alleles had comparatively low mean relative expression, while the more highly expressed alleles tended to have lower frequencies. This pattern was less evident at the supertype level (Fig. 2c, d), although the three supertypes with lowest mean relative expression (ST1: 0.067, ST2: 0.001, ST7: 0.0002) occurred in more than 75% of all individuals. Applying the same criteria of commonness applied for alleles (i.e. present in at least 5% of individuals), ST14 (overall 4 alleles of which 2 occur in the individuals included in this analysis, and mean relative expression = 1.2) was rare while all other supertypes were common (Fig. 2c, Table S3).

Similarly to the Midas cichlid, we found variation in MHC IIB expression in African cichlids (Kruskal-Wallis H = 311.96, df = 8, p < 0.001; Fig. 5a, Table S4) and these differences were consistent between species and sex (Fig. S1). Two putative loci (DBB & DCB) had low expression while the others had moderate to high expression. These two putative loci clustered with two Midas cichlid supertypes with exclusively lowly expressed alleles (DBB with ST1 and DCB with ST7), while most moderately to highly expressed putative loci clustered separately from Midas cichlid alleles (Fig. 3). While expression varied considerably among putative MHC IIB loci only one putative MHC IIA locus showed high expression, although all four differed significantly from each other (Kruskal-Wallis H = 252.47, df = 3, p < 0.001; Fig. 5b, Table S4).

**Figure 5.**
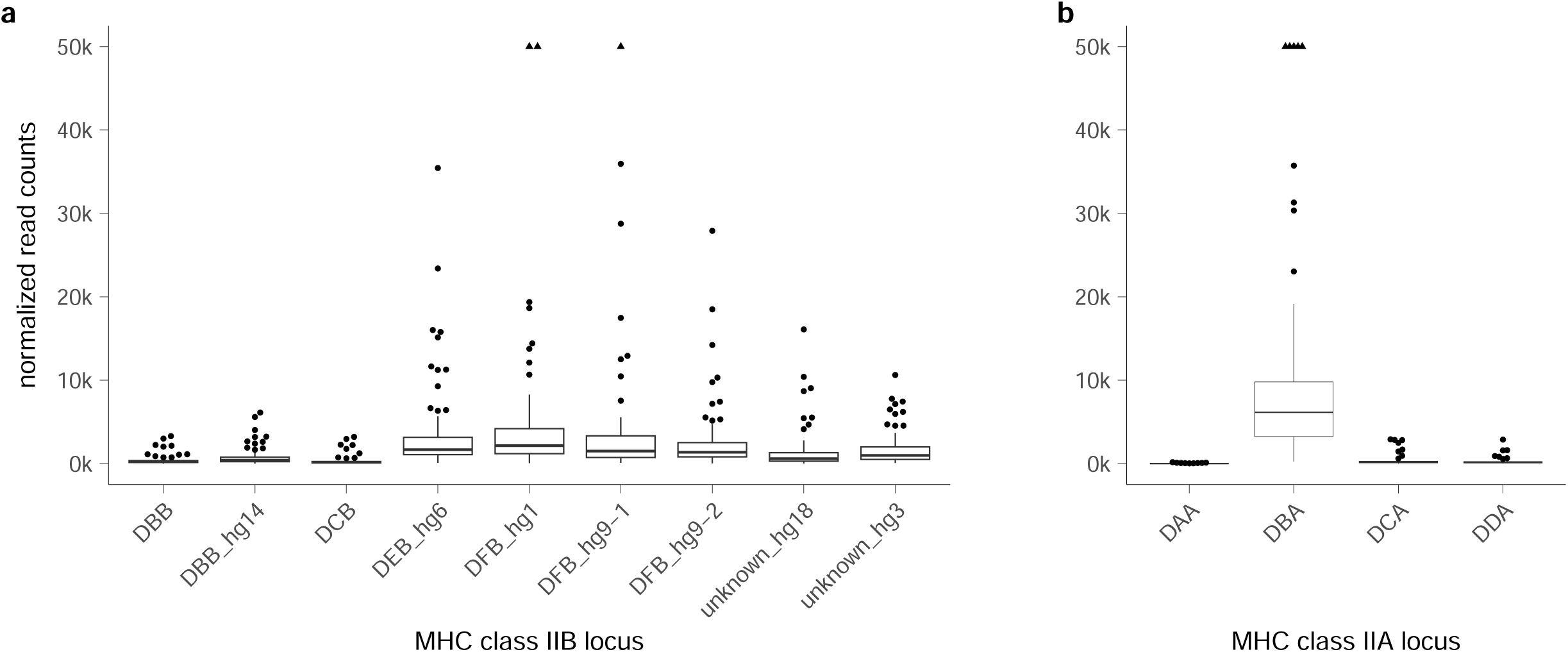
Expression of the putative loci of African cichlid MHC IIB (a) and IIA (b). Expression is approximated by normalized read counts (scaled by 1000). Read counts > 50000 are indicated as triangles.

For the Midas cichlid, the median number of alleles per individual was seven, and the median number of expressed alleles per individual was six (Table 2), and these two values were strongly correlated (r = 0.80, p < 0.01; Fig. 6a). When considering only *highly expressed* alleles (median of 4 alleles per individual), the correlation between number of alleles and number of expressed alleles per individual was even stronger (r = 0.98, p < 0.01; Fig. 6b). Only six individuals did not show expression of all alleles that were classified as *highly expressed*. Individuals possessed a median of three *lowly expressed* alleles of which a median of two were expressed. The ubiquitous *lowly expressed* allele Amci-DXB*040101 was always expressed. While individuals with more *lowly expressed* alleles also expressed more of these alleles, 259 individuals did not express all of the alleles classified as *lowly expressed* (r = 0.49, p < 0.01; Fig. 6c). The mean ± SD relative expression per individual was 2.3 ± 2.1 (*highly expressed*: 4.6 ± 4.1, mean ± SD *lowly expressed*: 0.02 ± 0.02; Table 2). Mean relative expression did not correlate significantly with numbers of alleles, although individuals with higher numbers of alleles tended to have lower mean expression (r = -0.11, p = 0.06; Fig. 6d). For *highly expressed* alleles, mean expression correlated negatively with the number of alleles (r = -0.21, p < 0.01; Fig. 6e) and more so for *lowly expressed* alleles (r = -0.38, p < 0.01; Fig. 6f). Larger deviations from the median number of *highly expressed* alleles were correlated with higher mean expression, although this correlation was not very strong (r = 0.13, p = 0.02). When considering all alleles or only *lowly expressed* alleles, there was no correlation between mean expression and numbers of alleles deviating from the median (r_all_ = 0.08, p_all_ = 0.18; r_low_ = -0.002, p_low_ = 0.98).

**Figure 6.**
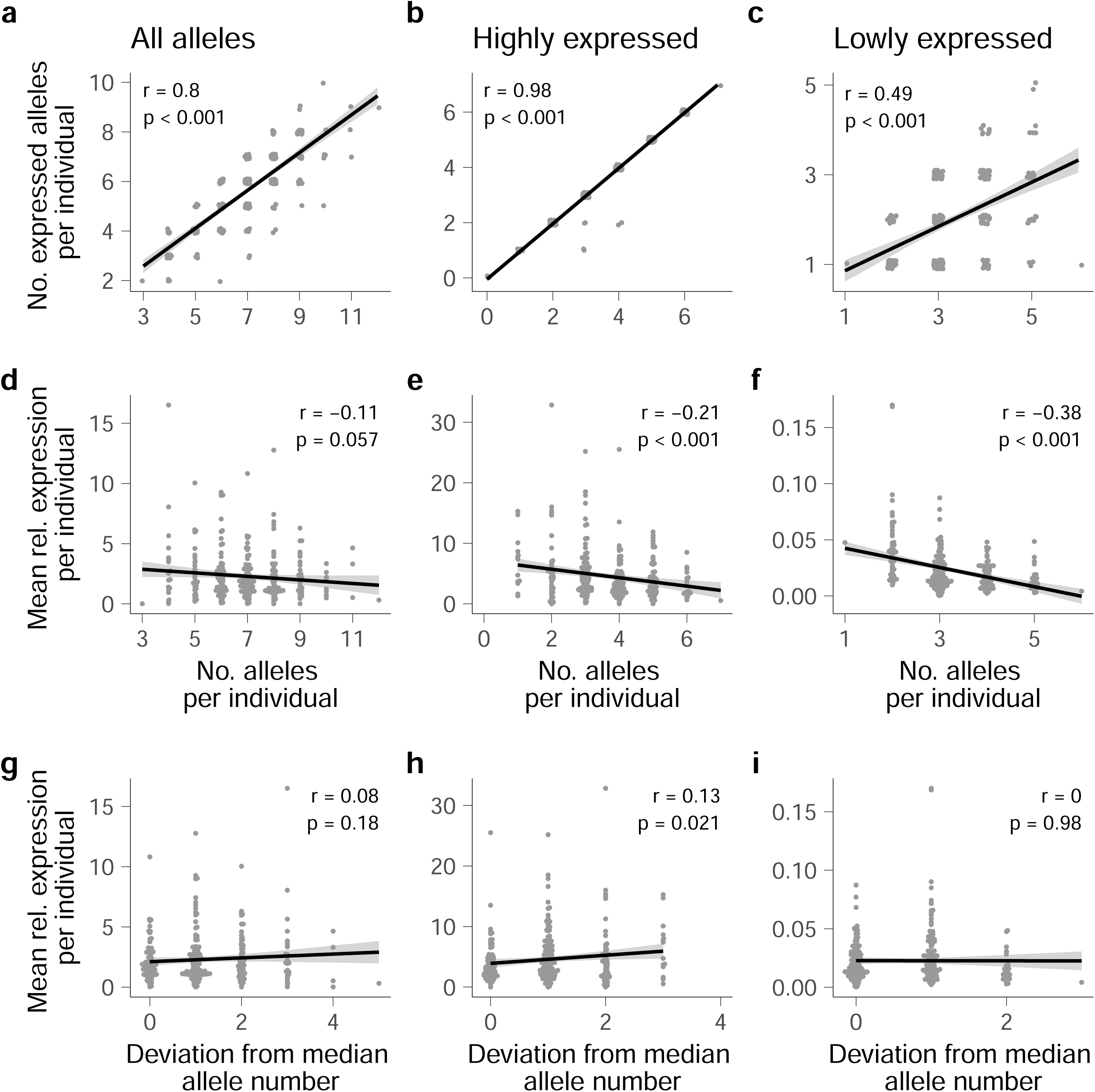
Correlation among numbers of alleles and relative expression per individual. Number of alleles in the genome *vs.* number of expressed alleles (a-c) and number of alleles *vs*. mean expression (d-f) for all alleles (a, d), only highly expressed alleles (b, e), and only lowly expressed alleles (c, f). Pearson’s r and p-values are given.

**Table 2.** Diversity statistics per individual.

| Row | Mean | SD | Median | Min | Max |
| --- | --- | --- | --- | --- | --- |
| No. alleles | 6.9 | 1.5 | 7 | 3 | 12 |
| No. highly expressed | 3.6 | 1.2 | 4 | 0 | 7 |
| No. lowly expressed | 3.3 | 0.88 | 3 | 1 | 6 |
| No. expressed alleles | 5.6 | 1.4 | 6 | 2 | 10 |
| Highly expressed | 3.6 | 1.3 | 4 | 0 | 7 |
| Lowly expressed | 2.0 | 0.88 | 2 | 1 | 5 |
| $\pi$ | 0.25 | 0.01 | 0.25 | 0.21 | 0.29 |
| $\pi$ expressed alleles | 0.24 | 0.02 | 0.24 | 0.16 | 0.29 |
| $\pi$ highly expressed alleles | 0.23 | 0.07 | 0.24 | 0.0 | 0.34 |
| $\pi$ lowly expressed alleles | 0.25 | 0.04 | 0.26 | 0.0 | 0.3 |
| Mean expression | 2.3 | 2.1 | 1.8 | 0.01 | 16.5 |
| Mean highly expressed | 4.6 | 4.1 | 3.5 | 0.0 | 32.8 |
| Mean lowly expressed | 0.02 | 0.02 | 0.02 | 0.0 | 0.17 |
| No. supertypes | 5.5 | 1.1 | 6 | 3 | 8 |
| No. expressed supertypes | 4.6 | 1.2 | 5 | 2 | 8 |

$\pi$ , nucleotide p-distance

Individuals possessed a median of six supertypes of which a median of five were expressed (Table 2). The majority of individuals carried only one allele of a given supertype, although up to three alleles of the same supertype were observed in single individuals. When combining the supertypes of exclusively *lowly expressed* alleles (ST1, ST2, and ST7), the median number of alleles per individual was three (range 1-6). However, not every individual possessed alleles of each *lowly expressed* supertype: ST1 was present in every individual due to the ubiquitous allele Amci-DXB*040101. Additionally, 239 individuals out of the 309 had at least one allele of each of ST2 and ST7, while one individual had only alleles of ST2 and 68 had only alleles of ST7. One individual did not possess any allele from ST2 or ST7.

## Discussion

This study presents an analysis of the expression levels of individual MHC alleles in a group of fish with one of the largest MHC class II allele repertoires described, the Neotropical Midas cichlid. In line with previous findings (Bracamonte et al., 2022), we found very large sequence diversity across the Midas cichlid populations, some common and ubiquitous alleles, and a large proportion of rarer alleles present in few individuals. Expression of alleles was highly variable, but the most abundant alleles showed low or no expression, while rare alleles showed the largest levels of expression. Therefore, rare alleles appear to be responsible for most resistance to parasites in this system.

Variation in expression levels among MHC alleles in non-model organisms was previously described for MHC class I (Drews et al., 2017; Drews & Westerdahl, 2019) and to some degree for MHC class II (Gaczorek et al., 2023; Schwensow et al., 2019). We found that expression of the MHC class II of the Midas cichlid varies considerably among alleles. Expression levels differ several orders of magnitude between *highly expressed* and *lowly expressed* alleles, and expression of alleles also differs among individuals. Also for the African MHC class II, we observed large expression differences among putative loci with variation both within and among species. Allele-specific expression has been shown to be modulated by genetic, epigenetic and environmental mechanisms (Handunnetthi et al., 2010; St. Pierre et al., 2022). Expression levels of MHC alleles and haplotypes were shown respond to individual infection states and environmental conditions (Axtner & Sommer, 2012), thus offering an additional level for selection to act upon. Despite that, we observed that MHC supertypes of alleles with putatively equivalent functionality of the Midas cichlid are composed of alleles of similar mean expression and that expression differences among loci of African cichlids were consistent among species. This suggests that, besides environmental factors, there may be a genetic component that regulates allele expression. For humans, variation in the promoter region was shown to affect expression levels of MHC class II alleles and even entire haplotypes (Handunnetthi et al., 2010). Why some alleles or haplotypes are generally more expressed than others is not fully understood for MHC class II. For the MHC class I, expression levels appear to be determined by the range of parasitic antigens that an MHC molecule can bind. Molecules that bind only few antigens have higher cell surface expression than alleles that bind a larger range of antigens (Chappell et al., 2015; Kaufman, 2018). Since interactions with parasitic antigens are similar between MHC class I and MHC class II (Morris et al., 1994), expression levels of MHC class II may also correlate with the range of antigens that an allele can respond to, which could explain why alleles within a supertype all show similar expression levels.

In the Midas cichlid MHC rare alleles were the most expressed and the most variable in expression, hence it is plausible that these alleles are interacting more tightly with parasites. Previous work in this system showed that many alleles are geographically restricted, only present in individuals from a single lake, or even specific to some populations, while groups of functional alleles or supertypes are generally widespread in lakes and populations (Bracamonte et al., 2022). If resistance relies on rare alleles, the capacity to respond to diverse parasite communities by different individuals and populations may be achieved with different alleles of the same supertypes.

We found clusters of lowly expressed alleles, similar to what was described in other systems for MHC class IIB and MHC class I (Drews et al., 2017; Drews & Westerdahl, 2019; Gaczorek et al., 2023). Two clusters of lowly expressed alleles (ST1 and ST2) had low sequence diversity and were not under positive selection, showing clear signatures of non-classical alleles (Adams & Luoma, 2013; Rodgers & Cook, 2005). Indeed, a previous study clustered these alleles more closely to another putative non-classical or pseudogene than to the remaining alleles (Hofmann et al., 2017). We observed a similar expression pattern for the African putative locus DBB, which clusters tightly with ST1, has a low level of polymorphism, and an amino acid conformation that may result in a different tertiary structure of the mature protein, again suggesting that it does not function as classical MHC (Hablützel et al., 2013). Non-classical alleles are common for both MHC class I and class II (Alfonso & Karlsson, 2000; Sato et al., 2001; Stet et al., 2003) and were previously associated with low expression levels (Drews et al., 2017). We found another cluster of lowly expressed alleles (ST7) with a less clear pattern. In this cluster, most alleles were not expressed and genetic diversity was low, compatible with a non-classic nature, but these alleles were under positive selection and phylogenetically similar to the putatively classical MHC alleles. Alleles of this supertype may have undergone recent pseudogenization with little time to disrupt the signal of positive selection. The putative African locus DCB showed a similar expression pattern to ST7, and was phylogenetically very close to this group of alleles. It is unlikely that the same set of alleles underwent pseudogenization independently in species that inhabit different continents and split 65 million years ago (Matschiner et al., 2020). Hence, these alleles may represent non-classical alleles or classical alleles with very restricted regulation of expression. Alternatively, they may be expressed in tissues not sampled for this study (Gerdol et al., 2019; Harstad et al., 2008) or at different time points during development.

At the individual level, we found that *highly expressed* alleles of the Midas cichlid were almost always expressed, regardless of how many such alleles an individual carried. This was not the case for *lowly expressed* alleles. Interestingly, individuals that possessed fewer alleles (both *highly expressed* and *lowly expressed*) had increased mean expression. Furthermore, deviation from the medium number of putatively classical alleles also led to increased mean expression. If we argue that the medium number of alleles per individual corresponds to the optimum number of alleles in the Midas cichlid, this deviation could indicate that expression levels compensate for possessing a subobtimal number of alleles. It has been argued that having a lower than optimal number of MHC alleles has an adverse effect on health due to the small number of parasites that can be detected (Woelfing et al., 2009) and increasing the levels of expression is a mechanism to mitigate this effect (Wegner et al., 2006). However, having a higher than optimal number of MHC alleles is also problematic due to the increased recognition of self-peptides which increases the risk of autoimmunity and reduces the number of T cells (Woelfing et al., 2009). Our results suggest that regulation of expression may also compensate for higher than optimal numbers of alleles. Fewer T cells may reduce the chance of interactions with T cells and elevated MHC expression may be a means of increasing the possibility that MHC binds to a T cell receptor upon encounter.

In conclusion, we found that expression levels differ markedly between common and less common alleles. Moderate to high expression was characteristic of less common alleles while low expression was predominantly found in the most common alleles. This suggests that rare MHC alleles could be more relevant for responding to parasite infections. Although the apparently relevant alleles were rare, the supertypes to which they belonged were widespread, thus maintaining resistance levels to parasites despite allele segregation or turnover. On the contrary, common alleles may have a different role in the immune response. Interestingly, the *lowly expressed* putative loci of the African cichlids clustered with the *lowly expressed* alleles of the Midas cichlid, which was not the case for most *highly expressed* alleles, providing further support that *lowly expressed* alleles/loci may have alternative functions. Overall, our results show that information on allele-specific or locus-specific expression of MHC in addition to sequence information can help identifying which alleles may be involved in parasite recognition. Furthermore, we provide evidence that commonness of an allele does not necessarily imply that it is relevant for defending against parasites.

## Acknowledgements

We thank Mariana Leal-Cardín, Ana Santacruz and the local fishermen in Nicaragua for their help with sample acquisition. We thank Dr. Elena Arriero, Jesús García, and Rosa María Pérez from the Universidad Complutense de Madrid for their help with the qPCRs and determining the optimal concentrations of blocking primers. We acknowledge the support of the technical personnel of the Laboratorio Sistemática Molecular y Genética de Poblaciones at MNCN for assistance with the laboratory procedures.

## Data Accessibility and Benefit-Sharing

### Data Accessibility

Raw sequencing data and related metadata of the Midas cichlid MHC will be made available as BioProjects upon acceptance. Midas cichlid MHC allele sequences will be submitted to NCBI upon acceptance.

### Benefit Sharing

Benefits Generated: Benefits from this research accrue from the sharing of our data and results on public databases as described above.

## Author Contributions

MB and SEB conceived the idea and designed the research. PIH contributed with African data. SEB and CLM did the lab work and SEB, PIH, and CLM analysed the data. SEB wrote the first draft and all authors contributed to the final version. All authors read and approved the final manuscript.

