## SupplementaryMaterial for "Expression diversity of cichlid MHC alleles"

### Table of Contents:


| | |
|---|---|
| **Table S1** | Page 2 |
| **Table S2** | Page 5 |
| **Table S3** | Page 10 |
| **Table S4** | Page 11 |
| **Figure S1** | Page 13 |

**Table S1.** African cichlid species used for the analysis

| Species | Sex | Accession number |
|---|---|---|
| Ctenochromis horei | female | SRR9680176 |
| Ctenochromis horei | female | SRR9680179 |
| Ctenochromis horei | female | SRR9680202 |
| Ctenochromis horei | male | SRR9680185 |
| Ctenochromis horei | male | SRR9680189 |
| Ctenochromis horei | male | SRR9680194 |
| Gnathochromis pfefferi | female | SRR9688133 |
| Gnathochromis pfefferi | female | SRR9688138 |
| Gnathochromis pfefferi | female | SRR9688475 |
| Gnathochromis pfefferi | male | SRR9688101 |
| Gnathochromis pfefferi | male | SRR9688571 |
| Gnathochromis pfefferi | male | SRR9688579 |
| Interochromis loocki | female | SRR9688507 |
| Interochromis loocki | female | SRR9688586 |
| Interochromis loocki | male | SRR9688516 |
| Interochromis loocki | male | SRR9688568 |
| Interochromis loocki | male | SRR9688590 |
| Interochromis loocki | male | SRR9688593 |
| Lobochilotes labiatus | female | SRR9686562 |
| Lobochilotes labiatus | female | SRR9686682 |
| Lobochilotes labiatus | female | SRR9686686 |
| Lobochilotes labiatus | male | SRR9686565 |
| Lobochilotes labiatus | male | SRR9686694 |
| Lobochilotes labiatus | male | SRR9686701 |
| Petrochromis famula | female | SRR9686397 |
| Petrochromis famula | female | SRR9686424 |
| Petrochromis famula | female | SRR9686493 |
| Petrochromis famula | male | SRR9686427 |
| Petrochromis famula | male | SRR9686467 |
| Petrochromis famula | male | SRR9686500 |
| Petrochromis fasciolatus | female | SRR9686513 |
| Petrochromis fasciolatus | female | SRR9686516 |
| Petrochromis fasciolatus | female | SRR9688403 |
| Petrochromis fasciolatus | male | SRR9688324 |
| Petrochromis fasciolatus | male | SRR9688402 |
| Petrochromis macrognathus | female | SRR9688255 |
| Petrochromis macrognathus | female | SRR9688281 |

| | | |
|---|---|---|
| Petrochromis macrognathus | female | SRR9688412 |
| Petrochromis macrognathus | male | SRR9688258 |
| Petrochromis macrognathus | male | SRR9688272 |
| Petrochromis macrognathus | male | SRR9688276 |
| Petrochromis polyodon | female | SRR9688230 |
| Petrochromis polyodon | female | SRR9688233 |
| Petrochromis polyodon | female | SRR9688409 |
| Petrochromis polyodon | male | SRR9688361 |
| Petrochromis polyodon | male | SRR9688366 |
| Petrochromis polyodon | male | SRR9688382 |
| Petrochromis trewavasae ephippium | female | SRR9686407 |
| Petrochromis trewavasae ephippium | female | SRR9686449 |
| Petrochromis trewavasae ephippium | female | SRR9686454 |
| Petrochromis trewavasae ephippium | male | SRR9686356 |
| Petrochromis trewavasae ephippium | male | SRR9686361 |
| Petrochromis trewavasae ephippium | male | SRR9686402 |
| Pseudosimochromis babaulti | female | SRR9687950 |
| Pseudosimochromis babaulti | female | SRR9687971 |
| Pseudosimochromis babaulti | female | SRR9688003 |
| Pseudosimochromis babaulti | female | SRR9688390 |
| Pseudosimochromis babaulti | female | SRR9688424 |
| Pseudosimochromis babaulti | female | SRR9688427 |
| Pseudosimochromis babaulti | male | SRR9687884 |
| Pseudosimochromis babaulti | male | SRR9687972 |
| Pseudosimochromis babaulti | male | SRR9688242 |
| Pseudosimochromis babaulti | male | SRR9688268 |
| Pseudosimochromis babaulti | male | SRR9688317 |
| Simochromis diagramma | female | SRR9688316 |
| Simochromis diagramma | female | SRR9688353 |
| Simochromis diagramma | female | SRR9688356 |
| Simochromis diagramma | male | SRR9688371 |
| Simochromis diagramma | male | SRR9688374 |
| Simochromis diagramma | male | SRR9688392 |
| Tropheus moorii | female | SRR9689088 |
| Tropheus moorii | female | SRR9689117 |
| Tropheus moorii | female | SRR9689233 |
| Tropheus moorii | male | SRR9689085 |
| Tropheus moorii | male | SRR9689122 |
| Tropheus moorii | male | SRR9689140 |
| Tropheus sp. black | female | SRR9689188 |
| Tropheus sp. black | female | SRR9689191 |

| | | |
|---|---|---|
| Tropheus sp. black | female | SRR9689236 |
| Tropheus sp. black | male | SRR9689237 |
| Tropheus sp. black | male | SRR9689261 |
| Tropheus sp. black | male | SRR9689262 |

**Table S2.** List of MHC alleles. The number of individuals in which it occurred (n), the functional supertype an allele was assigned to (Supertype), mean expression across individuals (Mean), standard deviation (SD), minimum expression (Min), and maximum expression (Max). Lowly expressed alleles are shaded in grey.

| Allele | n | Supertype | Mean | SD | Min | Max |
|---|---|---|---|---|---|---|
| ac151 | 1 | ST3 | 24.2 | | 24.2 | 24.2 |
| ac111 | 1 | ST3 | 14.2 | | 14.2 | 14.2 |
| ac161 | 1 | ST6 | 13.3 | | 13.3 | 13.3 |
| ac095 | 1 | ST11 | 13.2 | | 13.2 | 13.2 |
| ac039 | 8 | ST11 | 12.6 | 9.0 | 4.3 | 32.5 |
| ac005 | 26 | ST3 | 12.5 | 13.6 | 0.00031 | 62.5 |
| ac143 | 1 | ussigned 2 | 12.1 | | 12.1 | 12.1 |
| ac072 | 14 | ST3 | 11.7 | 12.5 | 0.0012 | 49.2 |
| Amci-DXB*2401 | 58 | ST6 | 11.3 | 9.5 | 0.0015 | 54.7 |
| ac082 | 2 | ST8 | 10.9 | 1.6 | 9.8 | 12.1 |
| ac028 | 4 | ST3 | 10.6 | 5.5 | 4.5 | 17.2 |
| ac056 | 1 | ST6 | 10.4 | | 10.4 | 10.4 |
| ac058 | 4 | ST11 | 10.1 | 6.0 | 4.6 | 18.7 |
| ac119 | 5 | ST6 | 9.8 | 7.4 | 1.5 | 19.3 |
| ac013 | 5 | ST4 | 9.8 | 1.5 | 7.8 | 12.0 |
| ac059 | 2 | ST3 | 9.5 | 3.5 | 7.1 | 12.0 |
| ac118 | 3 | ST6 | 9.3 | 5.1 | 3.6 | 13.5 |
| ac024 | 24 | ST4 | 8.6 | 6.8 | 1.7 | 32.4 |
| ac068 | 6 | ST10 | 8.6 | 11.7 | 0.0041 | 28.9 |
| ac074 | 7 | ST4 | 8.6 | 7.8 | 2.2 | 24.1 |
| ac011 | 5 | ST3 | 8.5 | 1.0 | 7.3 | 9.5 |
| ac090 | 1 | ST6 | 8.4 | | 8.4 | 8.4 |
| ac112 | 4 | ST11 | 8.3 | 3.9 | 4.0 | 13.4 |
| ac015 | 3 | ST3 | 8.3 | 8.9 | 0.0039 | 17.7 |
| Amci-DXB*09 | 9 | ST4 | 8.0 | 5.5 | 1.2 | 19.9 |
| ac033 | 2 | ST6 | 7.8 | 1.9 | 6.5 | 9.2 |
| ac047 | 2 | ST11 | 7.7 | 4.1 | 4.8 | 10.6 |
| ac008 | 10 | ST3 | 7.4 | 8.7 | 0.00023 | 27.7 |
| ac045 | 3 | ST3 | 7.4 | 6.2 | 1.7 | 14.0 |
| Amci-DXB*1001 | 26 | ST12 | 7.2 | 6.4 | 0.4 | 23.1 |
| ac012 | 20 | ST4 | 7.1 | 5.7 | 0.11 | 22.7 |
| ac110 | 1 | ST3 | 7.1 | | 7.1 | 7.1 |
| ac021 | 7 | ST3 | 7.0 | 5.0 | 0.0 | 13.8 |
| ac003 | 11 | ST3 | 7.0 | 2.5 | 3.3 | 12.1 |

| | | | | | | |
|---|---|---|---|---|---|---|
| ac026 | 7 | ST3 | 6.9 | 5.1 | 3.2 | 18.1 |
| ac086 | 2 | ST11 | 6.8 | 4.8 | 3.4 | 10.2 |
| ac007 | 13 | ST3 | 6.7 | 4.8 | 0.0039 | 20.7 |
| ac165 | 1 | ST10 | 6.7 | | 6.7 | 6.7 |
| ac141 | 2 | ST11 | 6.2 | 3.5 | 3.7 | 8.6 |
| ac009 | 19 | ST4 | 6.1 | 4.5 | 0.0 | 15.1 |
| ac043 | 10 | ST6 | 5.8 | 2.6 | 2.5 | 11.2 |
| ac004 | 25 | ST11 | 5.8 | 4.7 | 1.0 | 24.1 |
| ac006 | 15 | ST4 | 5.7 | 4.6 | 0.025 | 13.6 |
| ac080 | 1 | ST11 | 5.6 | | 5.6 | 5.6 |
| ac117 | 1 | ST5 | 5.5 | | 5.5 | 5.5 |
| ac032 | 17 | ST3 | 5.5 | 4.0 | 1.3 | 15.3 |
| Amci-DXB*21 | 9 | ST11 | 5.4 | 4.1 | 0.88 | 13.4 |
| Amci-DXB*25 | 37 | ST6 | 5.4 | 3.3 | 0.77 | 14.7 |
| ac081 | 3 | ST12 | 5.2 | 4.0 | 0.61 | 7.5 |
| ac029 | 19 | ST11 | 5.0 | 3.8 | 1.1 | 17.1 |
| ac037 | 2 | ST4 | 4.9 | 1.7 | 3.7 | 6.2 |
| ac132 | 4 | ST3 | 4.8 | 1.4 | 2.9 | 6.0 |
| ac054 | 2 | ST11 | 4.8 | 0.65 | 4.4 | 5.3 |
| ac148 | 1 | ST13 | 4.6 | | 4.6 | 4.6 |
| ac060 | 4 | ST6 | 4.6 | 2.2 | 2.9 | 7.8 |
| ac149 | 1 | ST13 | 4.5 | | 4.5 | 4.5 |
| ac010 | 19 | ST5 | 4.4 | 4.0 | 0.78 | 17.2 |
| Amci-DXB*19 | 17 | ST8 | 4.1 | 2.1 | 0.89 | 7.5 |
| ac097 | 5 | ST5 | 3.9 | 5.7 | 0.66 | 13.8 |
| ac063 | 2 | ST11 | 3.8 | 3.2 | 1.6 | 6.1 |
| ac025 | 6 | ST8 | 3.8 | 3.9 | 0.86 | 9.7 |
| ac089 | 3 | ST6 | 3.7 | 1.6 | 2.3 | 5.4 |
| ac061 | 6 | ST6 | 3.6 | 1.0 | 2.2 | 5.2 |
| ac094 | 2 | ST10 | 3.5 | 3.0 | 1.4 | 5.7 |
| ac052 | 3 | ST4 | 3.5 | 0.5 | 3.2 | 4.1 |
| ac053 | 3 | ST3 | 3.4 | 2.2 | 0.83 | 5.0 |
| ac129 | 1 | ST6 | 3.4 | | 3.4 | 3.4 |
| ac041 | 16 | ST11 | 3.3 | 2.7 | 0.0 | 11.4 |
| ac158 | 1 | ST5 | 3.2 | | 3.2 | 3.2 |
| ac070 | 6 | ST3 | 3.2 | 2.2 | 1.1 | 6.6 |
| ac077 | 1 | ST4 | 3.2 | | 3.2 | 3.2 |
| ac017 | 9 | ST8 | 3.2 | 1.0 | 1.4 | 4.9 |
| ac018 | 1 | ST11 | 3.0 | | 3.0 | 3.0 |
| Amci-DXB*110101 | 8 | ST5 | 2.9 | 2.3 | 0.3 | 7.6 |
| ac062 | 2 | ST3 | 2.9 | 2.0 | 1.4 | 4.3 |

| | | | | | | |
|---|---|---|---|---|---|---|
| ac138 | 2 | ST6 | 2.8 | 0.44 | 2.5 | 3.2 |
| ac067 | 4 | ST5 | 2.8 | 2.5 | 1.0 | 6.4 |
| Amci-DXB*12 | 8 | ST5 | 2.8 | 1.5 | 0.63 | 4.7 |
| ac159 | 1 | ST11 | 2.8 | | 2.8 | 2.8 |
| ac113 | 3 | ST5 | 2.7 | 4.2 | 0.00017 | 7.5 |
| ac101 | 5 | ST5 | 2.5 | 2.0 | 0.93 | 5.9 |
| ac076 | 1 | ST11 | 2.5 | | 2.5 | 2.5 |
| ac134 | 1 | ST12 | 2.4 | | 2.4 | 2.4 |
| ac031 | 3 | ST11 | 2.4 | 2.1 | 0.69 | 4.7 |
| ac136 | 3 | ST6 | 2.4 | 1.4 | 1.1 | 3.9 |
| ac046 | 1 | ST11 | 2.3 | | 2.3 | 2.3 |
| ac107 | 1 | ST5 | 2.3 | | 2.3 | 2.3 |
| ac016 | 33 | ST11 | 2.2 | 2.2 | 0.00052 | 8.5 |
| ac075 | 3 | ST6 | 2.1 | 0.8 | 1.2 | 2.8 |
| ac100 | 4 | ST11 | 2.1 | 2.5 | 0.0 | 5.6 |
| ac164 | 1 | ST11 | 2.0 | | 2.0 | 2.0 |
| ac170 | 1 | ST13 | 2.0 | | 2.0 | 2.0 |
| ac105 | 4 | ST5 | 2.0 | 2.1 | 0.0 | 4.8 |
| ac103 | 1 | ST11 | 2.0 | | 2.0 | 2.0 |
| ac035 | 4 | ST12 | 1.9 | 1.6 | 0.00009 | 4.0 |
| ac019 | 14 | ST8 | 1.8 | 1.3 | 0.41 | 5.2 |
| ac108 | 1 | ST5 | 1.7 | | 1.7 | 1.7 |
| ac038 | 10 | ST5 | 1.5 | 0.87 | 0.42 | 3.3 |
| Amci-DXB*18 | 1 | ST5 | 1.4 | | 1.4 | 1.4 |
| ac163 | 1 | ST14 | 1.3 | | 1.3 | 1.3 |
| Amci-DXB*060101 | 117 | ST9 | 1.3 | 1.5 | 0.00043 | 11.0 |
| ac157 | 1 | ST5 | 1.2 | | 1.2 | 1.2 |
| ac069 | 6 | ST10 | 1.2 | 0.4 | 0.71 | 1.6 |
| ac160 | 1 | ST14 | 1.2 | | 1.2 | 1.2 |
| ac171 | 1 | ST6 | 1.1 | | 1.1 | 1.1 |
| ac140 | 3 | ST11 | 1.1 | 0.25 | 0.78 | 1.2 |
| ac002 | 40 | ST9 | 0.96 | 0.98 | 0.00017 | 5.7 |
| ac096 | 1 | ST11 | 0.96 | | 0.96 | 0.96 |
| ac099 | 1 | ST5 | 0.94 | | 0.94 | 0.94 |
| Amci-DXB*13 | 18 | ST5 | 0.86 | 0.9 | 0.0 | 3.6 |
| ac042 | 1 | ST8 | 0.83 | | 0.83 | 0.83 |
| ac071 | 1 | ST11 | 0.79 | | 0.79 | 0.79 |
| ac153 | 1 | ST3 | 0.77 | | 0.77 | 0.77 |
| ac034 | 2 | ST11 | 0.76 | 0.44 | 0.45 | 1.1 |
| ac093 | 1 | ST5 | 0.74 | | 0.74 | 0.74 |
| Amci-DXB*08 | 15 | ST3 | 0.74 | 1.8 | 0.00008 | 7.0 |

| | | | | | | |
|---|---|---|---|---|---|---|
| ac115 | 1 | ST11 | 0.71 | | 0.71 | 0.71 |
| Amci-DXB*0604 | 119 | ST9 | 0.71 | 0.83 | 0.0 | 5.5 |
| ac145 | 1 | ST11 | 0.69 | | 0.69 | 0.69 |
| ac027 | 11 | ST8 | 0.62 | 0.42 | 0.00008 | 1.4 |
| ac083 | 1 | ST6 | 0.49 | | 0.49 | 0.49 |
| ac030 | 3 | ST6 | 0.35 | 0.16 | 0.2 | 0.52 |
| ac023 | 12 | ST8 | 0.3 | 0.29 | 0.0 | 1.1 |
| Amci-DXB*16 | 17 | ST5 | 0.28 | 0.17 | 0.0 | 0.6 |
| Amci-DXB*17 | 1 | ST5 | 0.28 | | 0.28 | 0.28 |
| ac050 | 4 | ST8 | 0.22 | 0.17 | 0.033 | 0.41 |
| ac135 | 1 | ST5 | 0.21 | | 0.21 | 0.21 |
| ac139 | 3 | ST8 | 0.19 | 0.06 | 0.13 | 0.25 |
| ac022 | 18 | ST5 | 0.19 | 0.2 | 0.0 | 0.85 |
| Amci-DXB*26 | 13 | ST13 | 0.17 | 0.17 | 0.032 | 0.62 |
| ac064 | 3 | ST3 | 0.16 | 0.12 | 0.07 | 0.3 |
| Amci-DXB*040101 | 309 | ST1 | 0.067 | 0.048 | 0.00068 | 0.34 |
| ac040 | 1 | ST3 | 0.04 | | 0.04 | 0.04 |
| Amci-DXB*0202 | 35 | ST2 | 0.0016 | 0.0053 | 0.0 | 0.032 |
| Amci-DXB*0203 | 21 | ST2 | 0.0016 | 0.0038 | 0.0 | 0.018 |
| Amci-DXB*0101 | 125 | ST2 | 0.00084 | 0.0033 | 0.0 | 0.036 |
| ac146 | 1 | ST2 | 0.00056 | | 0.00056 | 0.0006 |
| ac142 | 5 | ST7 | 0.0005 | 0.001 | 0.0 | 0.0023 |
| Amci-DXB*03 | 144 | ST2 | 0.00044 | 0.001 | 0.0 | 0.01 |
| ac152 | 1 | ST11 | 0.00029 | | 0.00029 | 0.0003 |
| ac001 | 306 | ST7 | 0.00024 | 0.0017 | 0.0 | 0.028 |
| ac091 | 3 | ST2 | 0.00004 | 0.00007 | 0.0 | 0.0001 |
| ac055 | 6 | ST7 | 0.00004 | 0.00006 | 0.0 | 0.0001 |
| ac014 | 51 | ST7 | 0.0 | 0.0 | 0.0 | 0.0 |
| ac044 | 1 | ST10 | 0.0 | | 0.0 | 0.0 |
| ac048 | 3 | ST7 | 0.0 | 0.0 | 0.0 | 0.0 |
| ac116 | 1 | ST7 | 0.0 | | 0.0 | 0.0 |
| ac120 | 1 | ST7 | 0.0 | | 0.0 | 0.0 |
| ac154 | 2 | ST7 | 0.0 | 0.0 | 0.0 | 0.0 |
| ac155 | 2 | ST7 | 0.0 | 0.0 | 0.0 | 0.0 |
| ac156 | 1 | ST11 | 0.0 | | 0.0 | 0.0 |
| ac162 | 1 | ST7 | 0.0 | | 0.0 | 0.0 |
| ac166 | 1 | ST7 | 0.0 | | 0.0 | 0.0 |
| ac167 | 1 | ST7 | 0.0 | | 0.0 | 0.0 |

**Table S3.** List of supertypes. Number of alleles per supertype (N allele), number of individuals a supertype occurred in (n), mean expression across individuals (Mean), standard deviation (SD), minimum expression (Min), and maximum expression (Max). Lowly expressed alleles are shaded in grey.

| Supertype | N allele | n | Mean | SD | pi | dN/dS | p |
|---|---|---|---|---|---|---|---|
| ussigned 2 | 1 | 1 | 12.1 | | | | |
| ST3 | 30 | 129 | 9.1 | 9.6 | 0.07 | 1.9 | 0.083 |
| ST6 | 29 | 132 | 8.5 | 7.7 | 0.062 | 1.3 | 0.3 |
| ST4 | 12 | 98 | 7.7 | 6.6 | 0.052 | 2.4 | 0.036 |
| ST12 | 5 | 34 | 6.3 | 6.0 | 0.055 | 2.4 | 0.12 |
| ST11 | 38 | 137 | 5.1 | 5.3 | 0.079 | 1.8 | 0.097 |
| ST10 | 6 | 16 | 4.6 | 7.7 | 0.084 | 0.84 | 1.0 |
| ST8 | 13 | 75 | 2.4 | 2.7 | 0.033 | 3.1 | 0.057 |
| ST5 | 28 | 116 | 2.1 | 2.8 | 0.048 | 1.2 | 0.39 |
| ST14 | 4 | 2 | 1.2 | 0.11 | 0.046 | 4.3 | 0.007 |
| ST9 | 4 | 225 | 1.2 | 1.4 | 0.019 | 0.56 | 1.0 |
| ST13 | 7 | 16 | 0.83 | 1.5 | 0.073 | 3.3 | 0.005 |
| ST1 | 4 | 309 | 0.067 | 0.048 | 0.032 | 1.0 | 0.49 |
| ST2 | 6 | 240 | 0.001 | 0.005 | 0.026 | 0.74 | 1.0 |
| ST7 | 14 | 307 | 0.000 | 0.002 | 0.022 | 3.2 | 0.035 |
| ussigned 1 | 1 | 0 | | | | | |

**Table S4.** Pairwise Mann-Whitney U tests of differential expression of African MHC IIB and MHC IIA alleles.

| Gene | Allele 1 | Allele 2 | W | Adj. p-value |
|---|---|---|---|---|
| **MHC IIB** | | | | |
| | DBB_hg14 | DBB | 4608 | 0.000 |
| | DCB | DBB | 2582 | 0.013 |
| | DEB_hg6 | DBB | 6249 | 0.000 |
| | DFB_hg1 | DBB | 6295 | 0.000 |
| | DFB_hg9-1 | DBB | 6109 | 0.000 |
| | DFB_hg9-2 | DBB | 6020 | 0.000 |
| | unknown_hg18 | DBB | 5023 | 0.000 |
| | unknown_hg3 | DBB | 5584 | 0.000 |
| | DCB | DBB_hg14 | 1551 | 0.000 |
| | DEB_hg6 | DBB_hg14 | 5790 | 0.000 |
| | DFB_hg1 | DBB_hg14 | 5835 | 0.000 |
| | DFB_hg9-1 | DBB_hg14 | 5447 | 0.000 |
| | DFB_hg9-2 | DBB_hg14 | 5370 | 0.000 |
| | unknown_hg18 | DBB_hg14 | 3974 | 0.050 |
| | unknown_hg3 | DBB_hg14 | 4718 | 0.000 |
| | DEB_hg6 | DCB | 6337 | 0.000 |
| | DFB_hg1 | DCB | 6387 | 0.000 |
| | DFB_hg9-1 | DCB | 6257 | 0.000 |
| | DFB_hg9-2 | DCB | 6169 | 0.000 |
| | unknown_hg18 | DCB | 5444 | 0.000 |
| | unknown_hg3 | DCB | 5870 | 0.000 |
| | DFB_hg1 | DEB_hg6 | 3657 | 0.342 |
| | DFB_hg9-1 | DEB_hg6 | 2930 | 0.165 |
| | DFB_hg9-2 | DEB_hg6 | 2768 | 0.056 |
| | unknown_hg18 | DEB_hg6 | 1465 | 0.000 |
| | unknown_hg3 | DEB_hg6 | 2050 | 0.000 |
| | DFB_hg9-1 | DFB_hg1 | 2706 | 0.036 |
| | DFB_hg9-2 | DFB_hg1 | 2511 | 0.007 |
| | unknown_hg18 | DFB_hg1 | 1383 | 0.000 |
| | unknown_hg3 | DFB_hg1 | 1903 | 0.000 |
| | DFB_hg9-2 | DFB_hg9-1 | 3184 | 0.559 |
| | unknown_hg18 | DFB_hg9-1 | 1852 | 0.000 |
| | unknown_hg3 | DFB_hg9-1 | 2474 | 0.005 |
| | unknown_hg18 | DFB_hg9-2 | 1948 | 0.000 |
| | unknown_hg3 | DFB_hg9-2 | 2633 | 0.021 |

| | | | | |
|---|---|---|---|---|
| | unknown_hg3 | unknown_hg18 | 4069 | 0.024 |
| **MHC IIA** | | | | |
| | DBA | DAA | 6724 | 0.000 |
| | DCA | DAA | 6513 | 0.000 |
| | DDA | DAA | 6054 | 0.000 |
| | DCA | DBA | 80 | 0.000 |
| | DDA | DBA | 38 | 0.000 |
| | DDA | DCA | 2665 | 0.022 |

p-values were adjusted using false discovery rate, W, Mann-Whitney U test statistic

**Figure S1.** Expression of the putative loci of African cichlid MHC IIB (a, c) and IIA (b, d) by species (a, b) and sex (c, d). Expression is approximated by normalized read counts (scaled by 1000).

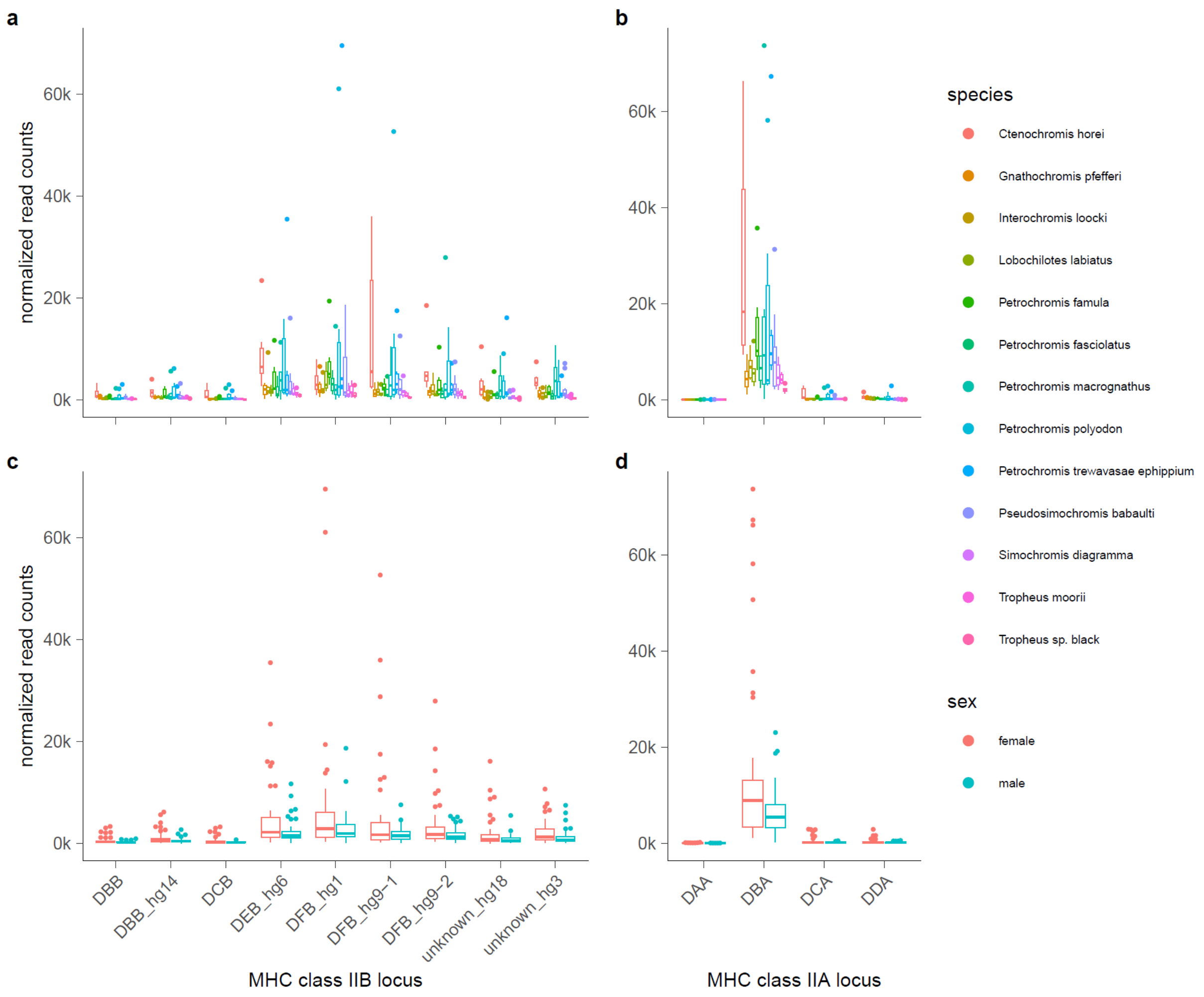
